# Loss of neurexins disrupts inhibitory connectivity and increases vulnerability of dopamine neurons in culture

**DOI:** 10.64898/2025.12.05.692551

**Authors:** Charles Ducrot, Alex Tchung, Samuel Burke, Consiglia Pacelli, Louis-Éric Trudeau

## Abstract

Midbrain dopamine (DA) neurons are essential regulators of basal ganglia function. Their axonal structure is intricate, with numerous non-synaptic release sites and fewer synaptic terminals that notably release glutamate or GABA. Despite their significance, the molecular mechanisms governing DA neuron connectivity and neurochemical identity remain poorly understood. We hypothesize that trans-synaptic cell adhesion molecules such as neurexins (Nrxns) regulate the interactions of DA neuron axons with target cells and thereby influence axonal branching and synapse formation by DA neurons. We therefore examined neuronal survival, axonal growth and synapse formation in cultured DA neurons lacking all neurexins (DAT::NrxnsKO). Conditional deletion of all Nrxns in DA neurons revealed that loss of Nrxns does not disrupt the basic development of these neurons or the structure of their axonal terminals, including normal expression of the vesicular monoamine transporter (VMAT2) and the calcium sensor synaptotagmin 1 (Syt1). However, loss of Nrxns affects the survival of DA neurons and their formation of inhibitory synapses, suggesting that Nrxns regulate the axonal connectivity of these neurons.

## Introduction

Ventral tegmental area (VTA) and substantia nigra compacta (SNc) dopamine (DA) neurons project densely to the ventral (vSTR) and dorsal striatum (dSTR), respectively, playing a critical role in modulating basal ganglia function (Descarries et al., 1980; Schultz, 2007; Matsuda et al., 2009). The axonal domain of these neurons is highly branched, containing a large number of non-synaptic release sites that release DA and a smaller subset of synaptic terminals that release glutamate and GABA (Dal Bo et al., 2004; Parent and Parent, 2006; Mendez et al., 2008; Stuber et al., 2010; Tritsch et al., 2012; Pacelli et al., 2015). Despite the well-established functional importance of DA in the brain, the molecular mechanisms governing the formation and regulation of release sites by DA neurons remain poorly (Mendez et al., 2011; Liu et al., 2018; Ducrot et al., 2021; Banerjee et al., 2022; Delignat-Lavaud et al., 2023).

Neurexins (Nrxns) are trans-synaptic proteins encoded by three genes that produce alpha (α) and beta (β) isoforms. They regulate the assembly, maturation, and function of neuronal synapses through interactions with postsynaptic partners such as neuroligins (NLs) (Ushkaryov et al., 1992; Ichtchenko et al., 1995; Graf et al., 2004; Chen et al., 2011; Südhof, 2017). While Nrxns are known to regulate fast synaptic neurotransmission, their role in regulating the formation and function of release sites established by neuromodulatory neurons such as DA neurons remains largely unexplored (Cheung et al., 2023; Ducrot et al., 2023). In previous work, we demonstrated that the conditional knockout (cKO) of Nrxns in DA neurons impairs DA signaling, as evidenced by slower DA reuptake, reduced density of the DA transporter (DAT), increased density of the vesicular monoamine transporter 2 (VMAT2), and decreased activity-dependent DA release. We also observed an increase of GABA co-transmission from DA terminals in the vSTR but not dSTR in these cKO mice, suggesting a region-specific regulatory role of Nrxns on GABA co-transmission in DA neurons (Ducrot et al., 2023).

Building on these findings, we aimed here to investigate whether these effects arise from deficits in axonal arborization and branching, and whether the proportion of DA neuron synapses capable of co-releasing both glutamate and GABA is altered. Given the difficulty of performing these experiments *in vivo*, we took advantage of an efficient *in vitro* model where midbrain DA neurons from the VTA or SNc were cultured with striatal neurons on an astrocyte monolayer. First, by combining fluorescence automated cell sorting (FACS) with RNA sequencing (RNAseq), we determined the molecular profile of DA neurons by examining the expression levels of synaptic genes, and compared this to GABAergic neurons, which are exclusively synaptic. We observed that the different isoforms of Nrxns are all expressed in DA neurons, but find that the expression levels vary between DA neurons in the VTA and SNc, suggesting a region-specific expression profile. We also found a relatively high expression of most exocytosis and active zone proteins, compared to striatal GABA neurons. To explore the role of Nrxns, we used a mouse model with conditional deletion of all three Nrxns genes in DA neurons (Ducrot et al., 2023) and assessed the axonal connectivity of DA neurons. Our findings revealed that while DA neurons cultured from DAT::Nrxns KO mice developed normally, their survival was reduced and the proportion of inhibitory synapses they established was diminished for both VTA and SNc neurons. Taken together, our results further strengthen the conclusion that the axonal connectivity of DA neurons is functionally regulated by Nrxns.

## Experimental procedures

### Animals

All procedures involving animals and their care were conducted in accordance with the Guide to care and use of Experimental Animals of the Canadian Council on Animal Care. The experimental protocols were approved by the animal ethics committees of the Université de Montréal (CDEA; #21-113). Housing was at a constant temperature (21°C) and humidity (60%), under a fixed 12h light/dark cycle with food and water available ad libitum.

### Generation of triple Neurexins cKO mice in DA neurons

All experiments were performed using mice generated by crossing DAT-IRES-Cre transgenic mice (Jackson Labs, B6.SJL-Slc6a3tm1.1 (Cre)Bkmn/J, strain 006660) with Nrxn123loxP mice [for details see (Ducrot et al., 2023; Brockhaus et al., 2024)]. Briefly, conditional knock-out (cKO) mice were produced as a result of CRE recombinase driving the selective excision of Nrxn-1, Nrxn-2 and Nrxn-3 genes in DA neurons and giving three different genotypes: Nrxn123 KO, Nrxn123 HET and Nrxn123 WT mice. The Nrxn123flox/flox mice were on a Cd1/BL6 mixed genetic background. The DAT-IRES-Cre mice were on a C57BL/6J genetic background. Male and female pups were used in all culture preparations.

### Genotyping

Mice were genotyped with a KAPA2G Fast HotStart DNA Polymerase kit (Kapa Biosystem). The following primers were used: DAT-IRES-Cre: Common 5’ TGGCTGTTGTGTAAAGTGG3’, wild-type reverse 5’GGACAGGGACATGGTTGACT 3’ and knock-out reverse 5’-CCAAAAGACGGCAATATGGT-3’, Nrxn-1 5’-GTAGCCTGTTTACTGCAGTTCATT-3’ and 5’-CAAGCACAGGATGTAATGGCCTT-3’, Nrxn-2 5’-CAGGGTAGGGTGTGGAATGAG-3’ and 5’-GTTGAGCCTCACATCCCATTT-3’, Nxn3 5’-CCACACTTACTTCTGTGGATTGC-3’ and 5’-CGTGGGGTATTTACGGATGAG-3’.

### Primary neuronal co-culture

For all experiments, postnatal day 0-3 (P0-P3) mice were cryoanesthetized, decapited and used for co-cultures according to a previously described protocol (Fasano et al., 2008; Ducrot et al., 2021). Primary VTA or SNc DA neurons were obtained from DAT::NrxnsKO or DAT::NrxnsWT pups and co-cultured with ventral striatum or dorsal striatum neurons obtained from DAT::NrxnsKO or DAT::NrxnsWT pups. Neurons were seeded on a monolayer of cortical astrocytes grown on collagen/poly-L-lysine-coated glass coverslips. All cultures were incubated at 37°C in 5% CO_2_ and maintained in 2/3 of Neurobasal medium, enriched with 1% penicillin/streptomycin, 1% Glutamax, 2% B-27 supplement and 5% fetal bovine serum (Invitrogen) plus 1/3 of minimum essential medium enriched with 1% penicillin/streptomycin, 1% Glutamax, 20mM glucose, 1mM sodium pyruvate and 100 µl of MITO+ serum extender. All primary neuronal co-cultures were used after 14 days *in vitro* (DIV14).

### Immunocytochemistry on cell cultures

Cultures were fixed at DIV14 with 4% paraformaldehyde (PFA; in PBS, pH-7.4), permeabilized with 0,1% triton X-100 during 20 min, and nonspecific binding sites were blocked with 10% bovine serum albumin during 10 min. Primary antibodies were: mouse anti-tyrosine hydroxylase (TH) (1:2000, Sigma), rabbit anti-TH (1:2000, Chemicon), rabbit anti-synaptotagmin 1 (Syt1) (1:1000, Synaptic Systems) and rabbit anti-vesicular monoamine transporter 2 (VMAT2) (1:1000, gift of Dr. Gary Miller, Colombia University). To improve immunoreactivity of the synaptic markers PSD95, gephyrin and bassoon, a set of cultures were fixed with 4% PFA together with 4% sucrose. For these experiments, the primary antibodies were mouse anti-PSD95 (1:1000 Pierce antibody), mouse anti-gephyrin (1:1000, Synaptic Systems) and guinea pig anti-bassoon (1:1000, Synaptic Systems). These were subsequently detected using Alexa Fluor-488-conjugated, Alexa Fluor-546-conjugated or Alexa Fluor-647-conjugated secondary antibodies (1:500, Invitrogen).

### RNA sequencing and data analysis

Transgenic mice expressing the enhanced green fluorescent protein (eGFP) gene in monoaminergic neurons under control of the TH promoter (TH-GFP mice) were used to isolate DA neurons and D2-GFP mice were used to isolated striatal GABA neurons. P0–P2 mice were cryo-anesthetized and decapitated for tissue collection. Freshly dissociated cells from the VTA or striatum were prepared as described previously (Mendez et al., 2008; Fulton et al., 2011), and GFP-positive neurons were purified by FACS and directly collected in Trizol (Qiagen). RNA extraction was performed with the RNeasy Mini kit (Qiagen) according to the manufacturer’s instructions. The concentration and purity of the RNA obtained from the neurons was determined using Qubit (Thermo Scientific) and quality was assessed with the 2100 Bioanalyzer (Agilent Technologies). Transcriptome libraries from three independent pools of VTA or SNc DA neurons, or from striatal GABA neurons were generated using the Truseq stranded mRNA library preparation kit (Illumina) with poly-A selection. Sequencing was performed on the HiSeq 2000 (Illumina), obtaining around 15M paired-end 100bp reads per sample. Statistical analysis was performed with DESeq software by using Wald-test as described previously (Love et al., 2014).

### Reverse transcription quantitative polymerase chain reaction (RT-qPCR)

We used RT-qPCR to quantify the amount of mRNA encoding the following genes: Nrxn-1α; Nrxn-1ß; Nrxn-1α; Nrxn-1ß; Nrxn-1α and Nrxn-1ß), from mesencephalic brain tissue from P0-P3 TH-GFP mice. The brains were quickly harvested and the ventral and dorsal striata were micro dissected and homogenized in 500 µL of trizol. RNA extraction was performed using RNAeasy Mini Kit (Qiagen, Canada) according to the manufacturer’s instructions. RNA integrity was validated using a Bioanalyzer 2100 (Agilent). Total RNA was treated with DNase and reverse transcribed using the Maxima First Strand cDNA synthesis kit with ds DNase (Thermo Fisher Scientific). For each qPCR assay, a standard curve was performed to ensure that the efficiency of the assay was between 90% and 110%. The primers used are listed in **supplementary Table 1**. Relative expression comparison (RQ = 2^^-ΔΔCT^) was calculated using GAPDH as an endogenous control.

### Image acquisition, neurite tracing and signal colocalization analysis

#### Image acquisition

All in vitro fluorescence imaging quantification analyses were performed on images acquired using an Olympus Fluoview FV1000 point-scanning confocal microscope (Olympus, Tokyo, Japan). Images were scanned sequentially to prevent non-specific bleed-through signal using 488, 546 and 647-nm laser excitation and a 60X (NA 1,42) oil immersion objective.

#### Cell body counts and neurite tracing

An Image-J (National Institute of Health, NIH) macro developed in-house was used to perform all quantifications. First, a background correction was applied for every image. Images were then segmented based on an intensity threshold to identify cell bodies. For validation, segmentation traces were then compared with raw images. They were then processed for tracing and quantification of neurites.

#### Colocalization analysis

Colocalization analysis was also performed using an in-house Image-J macro. First, the images were pre-processed by converting them to 8-bit, followed by enhancing the contrast and applying a slight blur to help segment axonal terminals. Following this step, a mask of axonal terminals was created by setting a threshold on the TH signal. This mask was then overlapped on top of the VMAT2, Syt1, bassoon or PSD-95 signal to quantify the proportion of TH-positive axonal terminals that are also positive for the other synaptic proteins. Secondly, we also performed a colocalization analysis using the JACoP (Bolte and Cordelières, 2006) plug-in to evaluate the Pearson’s coefficient between TH and the other synaptic proteins.

### Statistics

Data are presented as mean ± SEM. Statistically significant differences were assessed using a Student’s t-test for normally distributed data, or a non-parametric Mann-Whitney test for non-normally distributed data. For comparisons involving more than two groups with normal distributions, a two-way ANOVA followed by Sidak’s multiple comparison test was used (*p < 0.05; **p < 0.01; ***p < 0.001; ****p < 0.0001).

## Results

### Characterization of synaptic gene expression in FACS-purified DA neurons

Previous studies have provided evidence of Nrxn mRNA in mesencephalic DA neurons (Uchigashima et al., 2016, 2019), and we previously also provided evidence for expression of the three genes coding for Nrxn in DA neurons, but without distinguishing between the α- and β-isoforms (Ducrot et al., 2023). Moreover, very few studies have focused on mapping the molecular profile of exocytosis and active zone proteins in DA neurons. Here, we addressed this gap by quantifying the mRNA levels of synaptic proteins in postnatal VTA or SNc DA neurons isolated from transgenic mice that eGFP under control of the TH gene promoter (TH-GFP mice). For comparison, we also examined striatal neurons isolated from D2-GFP transgenic mice. We used FACS and RNAseq to analyze the expression level of each isoform of Nrxns in both DA neurons populations and in striatal neurons (**Fig. 1A** and **1B**). Our results show high abundance of mRNA for all three isoforms of Nrxn in both VTA and SNc DA neurons, with lower expression levels in striatal neurons. More specifically, Nrxn-1 was observed to be more expressed in VTA DA neurons, Nrxn-2 was similarly expressed in both regions, and Nrxn-3 was more abundant in SNc neurons (**Fig. 1C**). As a complementary analysis, we performed qPCR to compare the expression levels of α- and β-Nrxns isoforms in DA neurons of the VTA and SNc. We found that Nrxn-1α is expressed at approximately 2-fold higher levels in VTA DA neurons compared to SNc DA neurons. Additionally, we observed a trend toward higher expression of Nrxn-3β in SNc DA neurons compared to VTA DA neurons (**Fig. 1D** and **1E**). Examining the expression of other exocytosis-related genes more broadly, we find that all of these, including the exocytosis calcium-sensor Syt1, as well as several active zone proteins like Bassoon, RIM and ELKS, known to play a role in evoked DA release (Liu et al., 2018; Banerjee et al., 2020, 2022; Delignat-Lavaud et al., 2023), are expressed at higher levels in SNc DA neurons compared to VTA DA neurons or striatal neurons (**Fig. 1C**). This observation is in keeping with the previous demonstration that SNc DA neurons are endowed with larger axonal domains containing substantially more neurotransmitter release sites (Pacelli et al., 2015; Burke and Trudeau, 2022).

**Figure 1.**
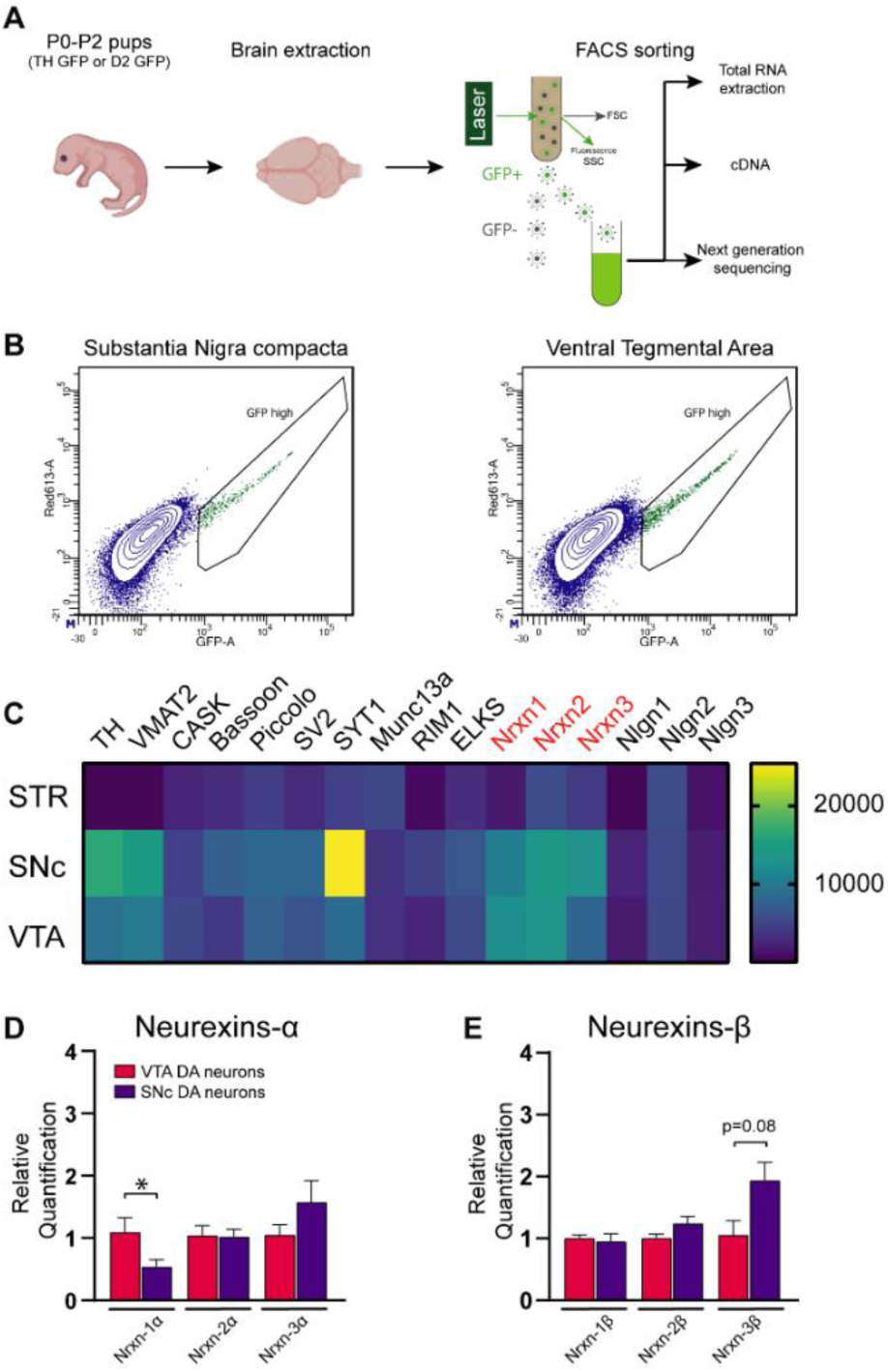
Relative abundance of mRNA encoding for neurexins and exocytosis-related genes in midbrain DA neurons compared to striatal GABA neurons. **A**-Schematic representation of the FACS procedure from P0-P2 TH-GFP and D2-GFP mouse pups. **B**-FACS analysis showing sorting of GFP-positive DA neurons from SNc (left) and VTA (right). **C** - RNAseq analysis to detect and quantify the relative mRNA levels of synaptic genes in postnatal (P0-P2) DA neurons or striatal neurons, with a heat map representing the relative expression of TH, VMAT2, CASK, Bassoon, Piccolo, SV2, Syt-1, Munc13, RIM, ELKS, Nrxn1, Nrxn2, Nrxn3, Nlgn1, Nlgn2, and Nlgn3. **D** and **E**-Relative quantification of mRNA encoding for α- and β-Nrxns obtained by RT-qPCR on VTA and SNc sorted DA neurons. Comparisons of data sets were made using an unpaired t-test, Nrxn-1α: VTA versus SNc, *p*<0.05. *The RNAseq values for neurexins, shown here for reference, were previously published in Ducrot et al., 2023. Similarly, the qPCR data for Nrxn1α are from Ducrot et al., 2021*.

### Dopamine neuron survival is decreased by deletion of all neurexins

In a previous study, we discovered that the absence of Nrxns in DA neurons disrupted GABAergic co-transmission specifically within DA neurons of the VTA, but not those of the SNc. One possible explanation is that the interaction between Nrxns and GABA_A_ receptors influences receptor localization and their activation by co-released GABA (Zhang et al., 2010). But loss of Nrxns could also potentially impact other aspects of the axonal connectivity of DA neurons. Previous data suggested that Nrxns can indeed influence the axonal development of neurons (Wang et al., 2019).We therefore examined the development of postnatal DA neurons *in vitro* as well as their basic connectivity. We deleted Nrxns 1, 2 and 3 from DA neurons by crossing Nrxn123^flox/flox^ mice with DAT-IRES-Cre mice (DAT::NrxnsKO) (**Fig. 2A**) and prepared primary co-cultures of SNc or VTA DA neurons together with ventral striatal neurons obtained from postnatal day 0-3 pups (P0-P3), as previously described (Ducrot et al., 2021). DA neurons were identified by TH immunocytochemistry and automated epifluorescence imaging (**Fig. 2B**). Intriguingly, the global survival of DA neurons in DAT::NrxnsKO cultures over 14 days was significantly reduced compared to WT controls (**Fig. 2C-E**) (two-way ANOVA, main effect of genotype; *p*<0.05). However, we did not detect genotype-specific changes in neurite development, quantified as the number and length of TH-positive neurite branches **(Fig. 2F and 2G**). A comparison of SNc and VTA cultures revealed a region-specific difference with slightly increased numbers of branches or branch length for VTA neurons (two-way ANOVA, main effect of region; p<0.05). We conclude that, although loss of Nrxns may decrease the basal resilience of DA neurons, it does not alter their intrinsic capacity to develop a complex axonal domain.

**Figure 2.**
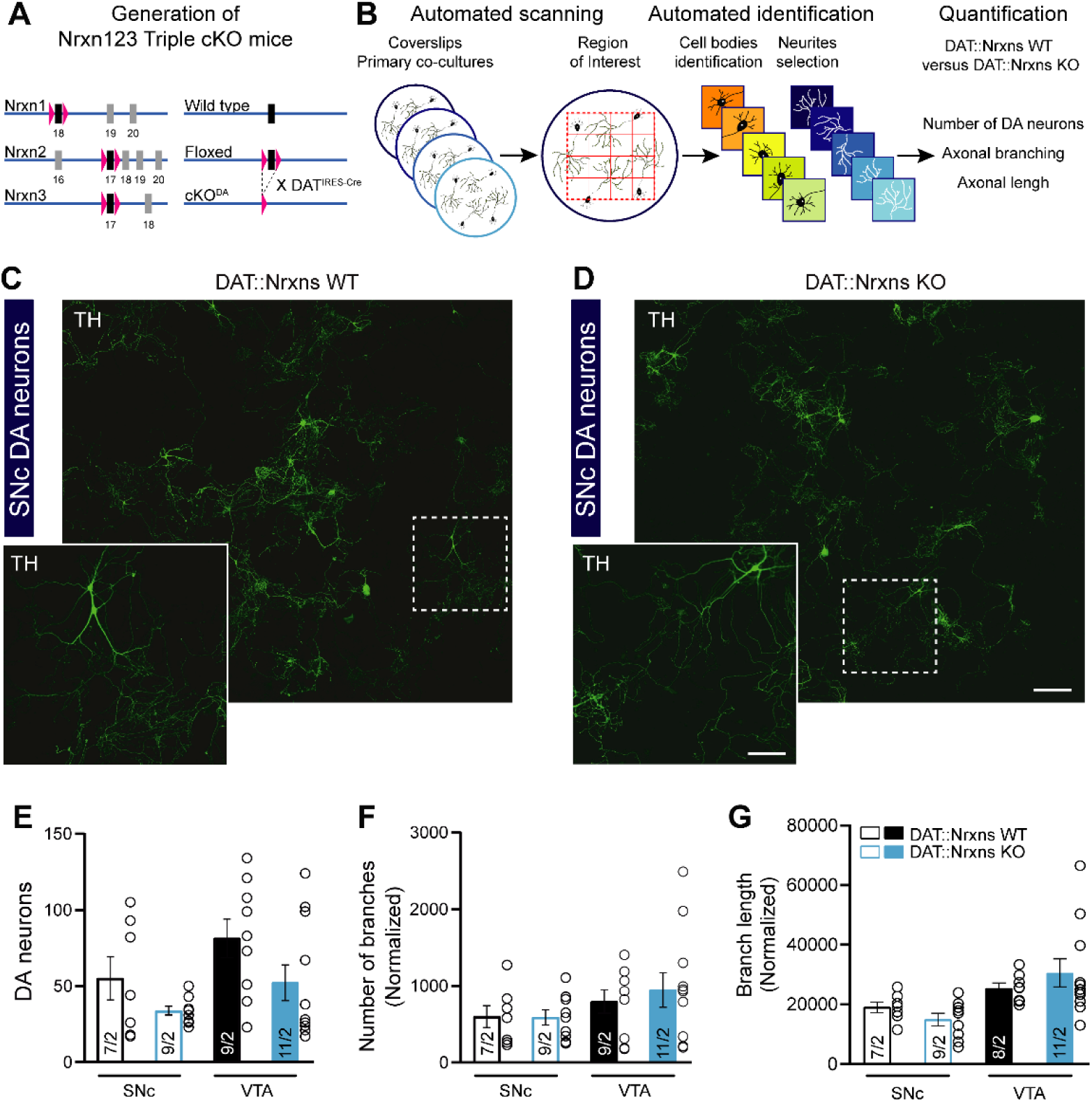
Normal axonal growth but reduced resilience of DA neurons after conditional deletion of all neurexins. **A**-Diagram explaining the genetic strategy for simultaneous conditional knockout of all neurexins in DA neurons **B** - Illustration of the method used to evaluate and quantify the survival and growth of DA neurons lacking all neurexins. **C** and **D**-Representative TH+ SNc DA neurons at DIV14 from DAT::NrxnsWT and DAT::NrxnsKO **E** - Quantification of DA neuron survival, assessed by their number per coverslip at DIV14 (SNc WT=55.00 ± 14.18; SNc KO=33.78 ± 2.84; VTA WT=81.33 ± 12.62; VTA KO=52.18 ± 11.72). The comparison was performed using a 2-way ANOVA followed by Šidák’s multiple comparison test. Main effect of genotype *p*<0.05. **F** and **G** - Evaluation of neurite development in cultured DA neurons, assessed by quantifying the number of TH-positive processes (**F**) and their length (**G**) (*Branch number*: SNc WT= 598.9 ± 143.3, SNc KO= 588.2 ± 99.37, VTA WT= 795.4 ± 152.6, VTA KO= 945.7 ± 225.7; *Branch length*: SNc WT= 19086 ± 1749µm, SNc KO= 14950 ± 2068 µm, VTA WT= 34721 ± 9282 µm, VTA KO= 30593 ± 4715 µm, Two-way ANOVA, *p*=0.47). For axonal growth assessment: n = 7-11 coverslips from 2 different neuronal co-cultures. For all analyses, the plots represent the mean ± SEM. Statistical analysis was carried out by a 2-way ANOVA followed by Šidák’s multiple comparison test.

### Similar expression of presynaptic markers in DAT::NrxnsWT and DAT::NrxnsKO DA neurons

Even if global axonal development appeared unimpaired, it was considered possible that loss of Nrxns would perturb the integrity of DA neuron axonal terminals. In a second set of analyses, we therefore examined some of the characteristics of axonal terminals established by DAT::NrxnsKO DA neurons. After double-labelling for TH and the ubiquitous exocytosis Ca^2+^ sensor Syt1, fields containing axonal arbors were examined for colocalization analysis (**Fig. 3A**, **3B**, **3C**, **3D**). We found that Syt1 and VMAT2 were detectable in a majority of TH-positive axonal terminals, in keeping with previous work (Ducrot et al., 2021). Although the effect size was modest, we found that the proportion of dopaminergic terminals containing Syt1 was higher in DA neurons lacking Nrxns (**Fig. 3E**, two-way ANOVA, main effect of genotype; *p* < 0.05). It was also higher in SNc DA neurons compared to VTA DA neurons (**Fig. 3E**, two-way ANOVA, main effect of region; *p* < 0.005). However, double labelling for TH and VMAT2, the transporter responsible for vesicular packaging of DA (**Fig. 3C** and **3D**), showed no significant change in the proportion of DA terminals containing VMAT2 (**Fig. 3F**, two-way ANOVA, *p* > 0.05). We conclude that loss of Nrxns does not impair the intrinsic capacity of these neurons to establish neurotransmitter release sites.

**Figure 3.**
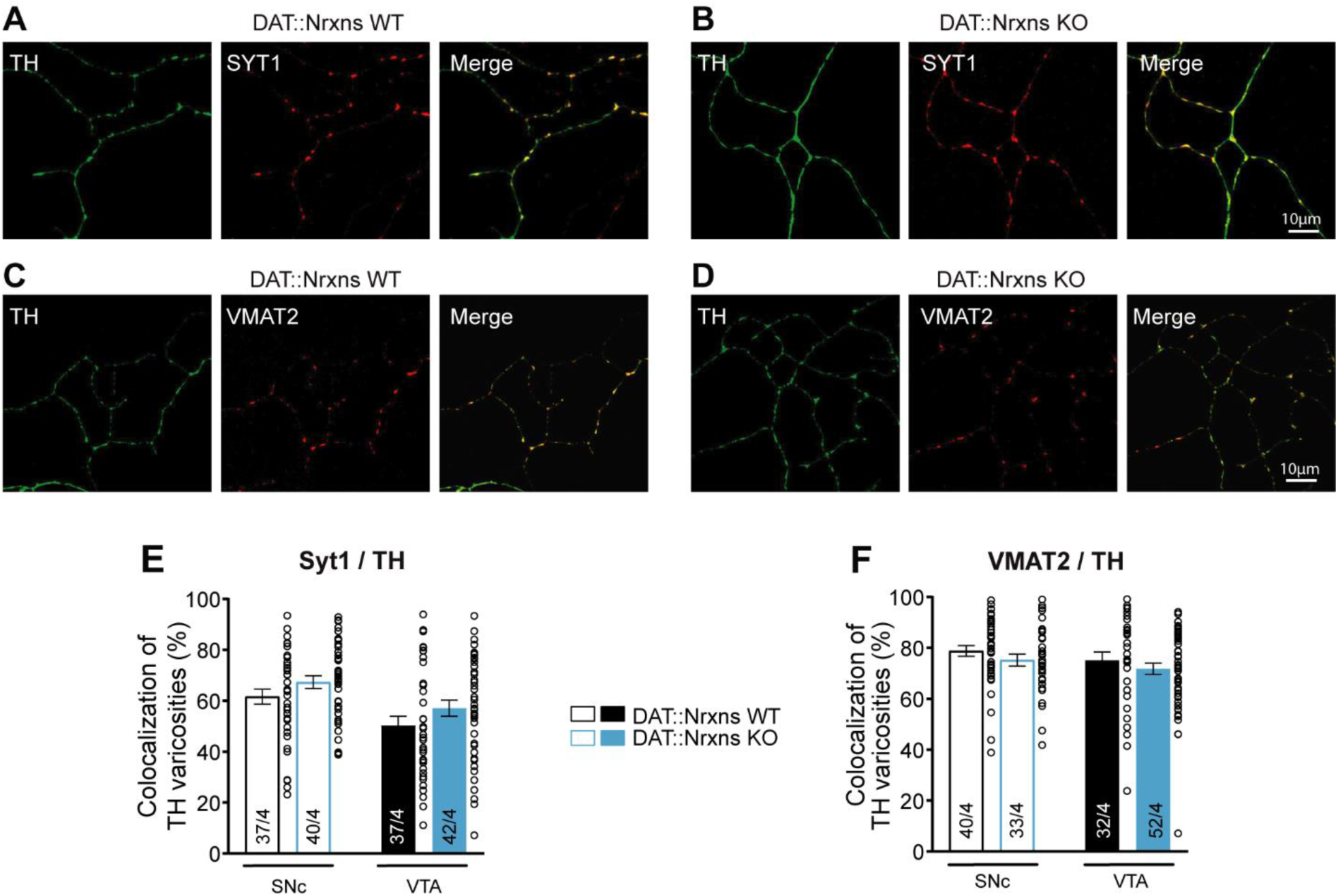
Normal expression of Syt1 and VMAT2 in the axonal terminals of DA neurons after conditional deletion of all neurexins. **A** and **B**-Photomicrographs illustrating the distribution of TH/Syt1 positive axonal terminals along the axonal domain of a SNc DA neuron from DAT::NrxnsWT (**A**) and DAT::NrxnsKO (**B**) mice. **C** and **D-** Photomicrographs illustrating the distribution of TH/VMAT2 positive axonal terminals along the axonal domain of a SNc DA neuron from DAT::NrxnsWT (**C**) and DAT::NrxnsKO (**D**) mice. **E-** Summary graph showing the proportion of TH-positive terminals containing Syt1. The comparison was carried out by a 2-way ANOVAs followed by Šidák’s multiple comparison test. Main effect of genotype *p* < 0.05 and region *p*<0.005. **F**-Summary graph showing the proportion of TH-positive terminals containing VMAT2. For Syt1/VMAT2 quantifications: n = 32-52 axonal fields from 4 different neuronal co-cultures. The number of observations represent the number of fields from individual neurons examined. For all analyses, the plots represent the mean ± SEM. Statistical analysis was carried out by 2-way ANOVAs followed by Šidák’s multiple comparison.

### Neurexin deficiency disrupts the proportion of inhibitory synapses established by DA neurons

A subset of terminals along the complex axonal arbor of DA neurons has the capacity to release glutamate or GABA (Sulzer et al., 1998; Dal Bo et al., 2004; Mendez et al., 2008; Stuber et al., 2010; Tritsch et al., 2012, 2016). We hypothesized that deletion of Nrxns could alter the formation of excitatory or inhibitory synapses by cultured SNc and VTA DA neurons. To test this, we co-cultured DA neurons with striatal neurons, and examined DA neuron axon terminals in close proximity to postsynaptic organizers associated with glutamate (PSD95) and GABA (gephyrin) synapses (Craig et al., 1996; Kornau et al., 1997). Loss of Nrxns led to a modest, but significant reduction in the proportion of DA neuron terminals colocalizing with gephyrin in DAT::NrxnsKO mice compared to controls, independent of the region examined (**Fig. 4A-C**; two-way ANOVA, main effect of genotype, *p* < 0.05). In contrast, the proportion of SNc and VTA DA terminals associated with PSD95 was unaffected by deletion of Nrxns (**Fig. 4D** and **Fig S1**; two-way ANOVA, *p* > 0.05). In previous work, we demonstrated that most Bassoon-positive DA neuron terminals are typically located near target cells (Ducrot et al., 2021; Bulumulla et al., 2024). Previous studies have also shown that in the absence of all Nrxns, the expression of bassoon is slightly but significantly reduced in the presynaptic active zone of the calyx of Held (Luo et al., 2020). Based on these findings, we evaluated the proportion of DA neuron terminals expressing the active zone protein Bassoon. We find that in the absence of Nrxns, there is no difference in the proportion of VTA and SNc DA terminals colocalizing with Bassoon (**Fig. 4E**; two-way ANOVA, *p* > 0.05). We conclude that these transsynaptic proteins play a limited role in regulating the organization of axonal terminals along the axonal arbor of DA neurons.

**Figure 4.**
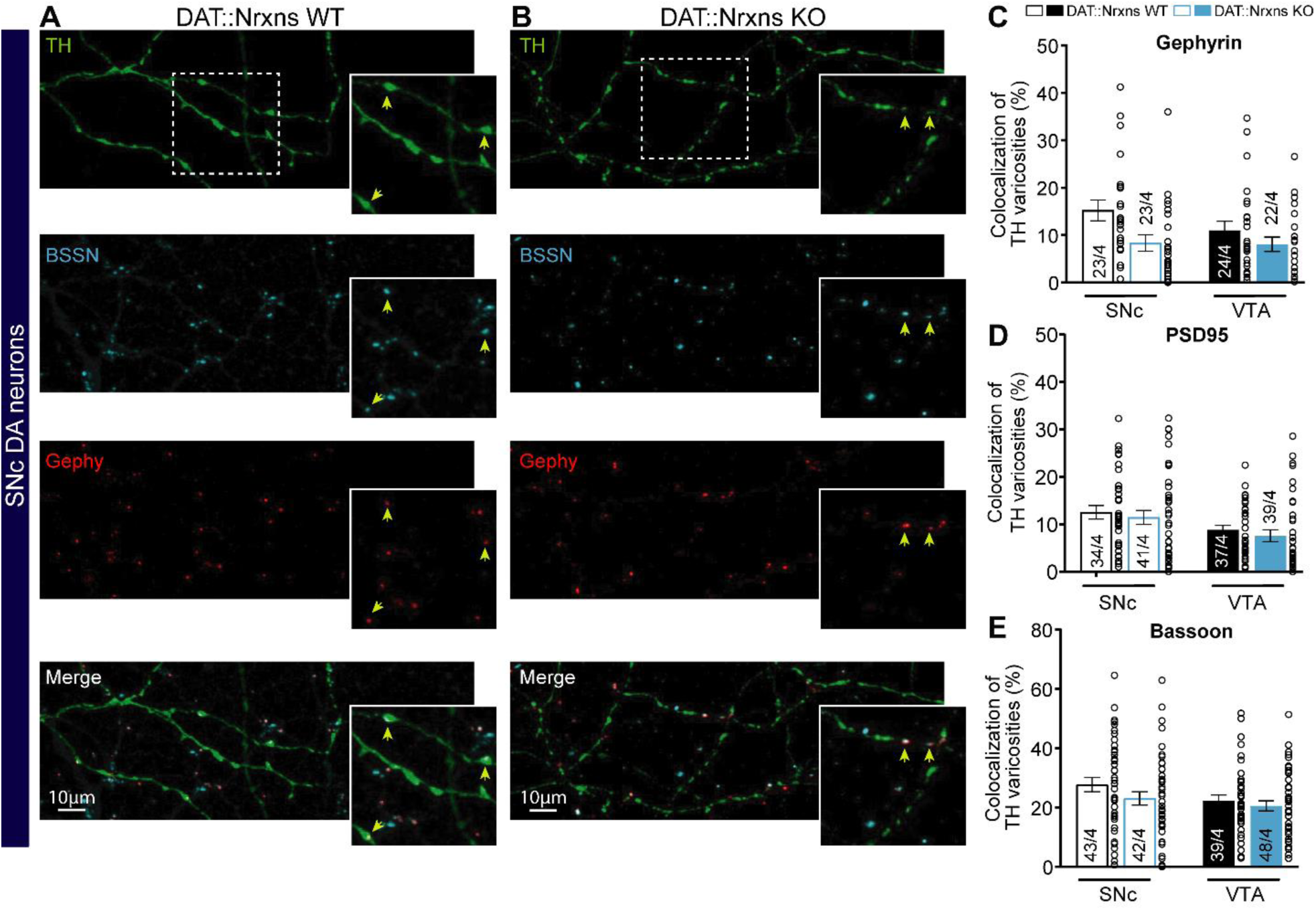
Reduced proportion of inhibitory synapses established by cultured DA neurons after conditional deletion of neurexins. **A** and **B-** Representative images illustrating TH-positive terminals in the axonal arbor of a SNc DA neuron from DAT::NrxnsWT (**A**) and DAT::NrxnsKO (**B**) mice. Bassoon was expressed sparsely and colocalized with the postsynaptic marker gephyrin. Yellow arrows indicate regions where TH, Bassoon, and Gephyrin colocalized. **C-** Bar graph representing the proportion (%) of axonal terminals established by VTA and SNc DA neurons that are positive for TH and colocalizing with gephyrin at DIV14. The comparison was carried out by a 2-way ANOVA followed by Šidák’s multiple comparison. Main effect of genotype *p*< 0.05. **D-** Bar graph representing the proportion (%) of axonal terminals established by VTA and SNc DA neurons that are positive for TH and colocalizing with PSD95 at DIV14. **E-** Bar graph representing the proportion (%) of axonal terminals that are positive for bassoon in VTA and SNc DA neurons from DAT::NrxnsWT and DAT::NrxnsKO mice. Data represent mean ± SEM. Statistical analysis was carried out by 2-way ANOVAs followed by Šidák’s multiple comparison.

## Discussion

Nrxns are key presynaptic organizers that couple Ca²⁺ channels to the release machinery and coordinate synapse differentiation through interactions with diverse postsynaptic partners (Ushkaryov et al., 1992; Missler et al., 2003; Südhof, 2017). While their role is well established in classical glutamatergic and GABAergic neurons, how Nrxns contribute to neuromodulatory systems such as DA neurons has remained largely unexplored.

We previously reported that deletion of Nrxns in DA neurons alters DA signaling and enhances GABA co-transmission in the ventral striatum (Ducrot et al., 2023). Building on this work, we now show that Nrxns also regulate axonal connectivity and synapse specification in midbrain DA neurons. We find that loss of all three Nrxns genes does not affect overall axonal architecture or the formation of release sites but causes a small reduction in DA neuron survival and selectively decreases the proportion of inhibitory, rather than excitatory, synaptic contacts formed by VTA and SNc neurons. These results indicate that Nrxns shape DA neuron connectivity, extending their established roles in fast synapses to neuromodulatory circuits.

To dissect how Nrxns influence DA neuron organization at the cellular level, we turned to primary cultures where individual presynaptic and postsynaptic components can be examined with high resolution. We found that deletion of Nrxns does not alter the expression or localization of core presynaptic markers such as VMAT2 and Syt1, indicating that the machinery for DA storage and calcium-dependent release remains intact. The modest increase in Syt1-positive terminals further suggests a compensatory reinforcement of release competence rather than a structural loss of terminals. These results reveal that DA neurons can still form functional release sites without Nrxns, consistent with their dual organization including many non-synaptic terminals for volume transmission and a smaller subset of true synaptic terminals for fast signaling. Overall, our findings indicate that Nrxns are dispensable for building the basic release machinery but contribute to defining synaptic identity, particularly at inhibitory sites, aligning with studies showing that Nrxns modulates the brain’s excitatory/inhibitory balance (Ullrich et al., 1995; Graf et al., 2004; Boucard et al., 2005; Chih et al., 2006; Zhang et al., 2010; Aoto et al., 2013; Südhof, 2017; Boxer and Aoto, 2022).

This selective influence on synaptic identity is further supported by our morphological analyses, showing that the triple KO of Nrxns in DA neurons led to a reduction in Gephyrin-positive contacts in culture, suggesting that Nrxns are required for the stabilization or maturation of inhibitory-like synapses. This structural deficit contrasts with our previous *in vivo* findings showing enhanced GABA co-release (Ducrot et al., 2023), implying that Nrxns exert distinct, context-dependent roles in DA neurons. While they appear essential for the adhesive organization of inhibitory contacts *in vitro*, Nrxns may also constrain GABAergic output in the intact brain, possibly through direct trans-synaptic signaling with GABA_A_ receptors. These data indicate that Nrxns exert both structural and functional control over inhibition, modulating rather than defining connectivity in the DA system.

Interestingly, the deletion of Nrxns affected not only synaptic organization but also DA neuron survival, revealing a potential trophic role beyond synapse assembly. While this phenotype could arise indirectly from impaired synaptic support or altered communication with target neurons, it is also possible that Nrxns contribute cell-autonomously to DA neuron resilience. Recent studies have shown that presynaptic adhesion molecules influence axonal growth and maintenance through intracellular signaling pathways mediated by the cytoplasmic domains of Nrxns (Dean et al., 2003; Wang et al., 2019). The loss of these interactions may compromise axonal stability or trophic signaling, ultimately reducing neuronal survival. Given that DA neurons are particularly vulnerable to metabolic and oxidative stress (Pacelli et al., 2015), even subtle disturbances in connectivity could have cumulative effects on viability, especially under *in vitro* conditions, that are known to be stressful for DA neurons. Interestingly, similar findings were reported *in vivo* in serotonergic neurons, where the deletion of Nrxns caused a significant loss of cell bodies in the dorsal and magnus raphe nuclei (Cheung et al., 2023). These results are consistent with our observations and support a conserved role for Nrxns in regulating neuronal survival. Although we did not detect changes in axonal arborization in our cultures, this discrepancy may reflect model-specific differences, as our *in vitro* system was maintained for only 14 days, whereas *in vivo* analyses were performed at 8 weeks. Despite these alterations in inhibitory connectivity and cell survival, other aspects of presynaptic organization remained largely unaffected.

Notably, we found no significant difference in Bassoon localization in DAT::NrxnsKO DA neurons, suggesting that Nrxns are not required for the recruitment of active zone scaffolds at DA terminals. This differs from observations at classical synapses such as the calyx of Held, where Nrxns deletion slightly reduced Bassoon clustering (Luo et al., 2020). Such differences may reflect the unique organization of DA neuron axons, where active zone-like structures are sparse and may form through alternative adhesion systems. The preservation of Bassoon expression in our cultures is consistent with the notion that DA neurons rely on multiple, partially redundant pathways to assemble presynaptic release machinery.

Our analysis of neurite complexity revealed an increased number of branches and branch length in VTA compared to SNc DA neurons. This was surprizing considering our previous work showing increased axonal volume in SNc compared to VTA DA neurons, both in vitro (Pacelli et al., 2015) and in vivo (Giguère et al., 2019). However, it should be noted that in the present study, we did not distinguish between axonal and dendritic branches due to limitations in antibody combinations. Also, the previous observation of increased axonal complexity in SNc compared to VTA DA neurons was observed after 3 and 7 days *in vitro*. It is possible that under *in vitro* conditions that lack the guidance and trophic signals found *in vivo*, initial intrinsic difference in axonal growth become lost over time.

Altogether, our results identify Nrxns as key regulators of DA neuron organization, acting to promote inhibitory synapse formation and sustain neuronal survival without altering the global axonal architecture or DA neuron release sites. This selective influence likely contributes to the balance between synaptic and non-synaptic transmission modes characteristic of DA neurons. Because the expression of Nrxns varies across DA neuron subpopulations, with distinct isoform combinations in VTA and SNc neurons (Uchigashima et al., 2019; Ducrot et al., 2023), future studies should determine whether specific isoforms confer region- or target-specific connectivity features. It will also be important to explore whether altered Nrxn signaling contributes to the vulnerability of DA neurons in pathological contexts such as Parkinson’s disease or addiction, where synaptic remodeling and axonal degeneration are prominent.

## Author contributions

C.D: Conceptualization; Data curation; Formal analysis; Methodology; Writing - original draft, Writing review & editing. A.T: Formal analysis, S. B-N: Formal analysis and Methodology, C.P: Methodology, L-E. T: Funding acquisition; Supervision; Writing review & editing.

## Acknowledgements

We would like to thank Marie-Josée Bourque for excellent technical assistance with neuronal cultures and for stimulating scientific discussions, Dr. G. Miller for kindly providing the VMAT2 antibody, the IRIC genomic platform for the sequencing analysis and Dr L.Y Chen (University of California, Irvine) for providing the Nrxn1/2/3 conditional knockout mouse line. This work was funded by Canadian Institutes of Health Research (CIHR, grant PJT-183931) and C.D. received a graduate student award from Fond de Recherche en Santé du Québec (FRQS, grant #255931).

---

**Supplemental table 1.**
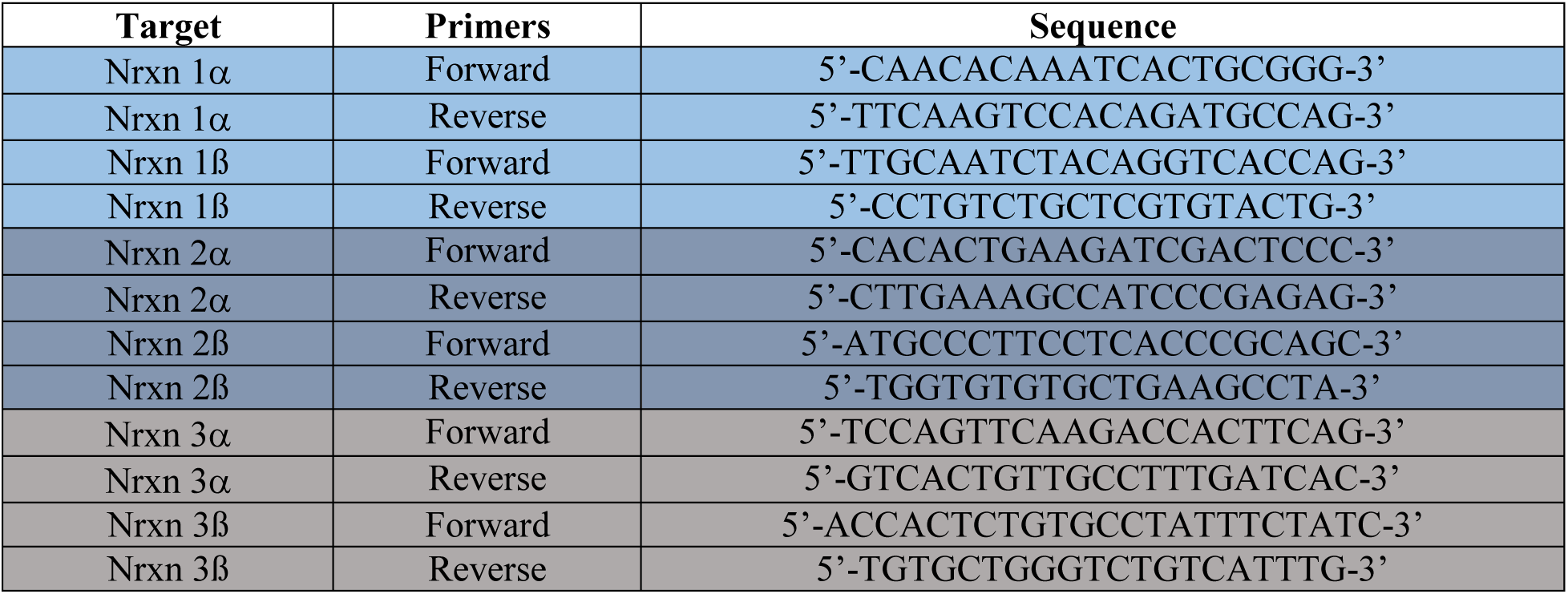

**Figure S1.**
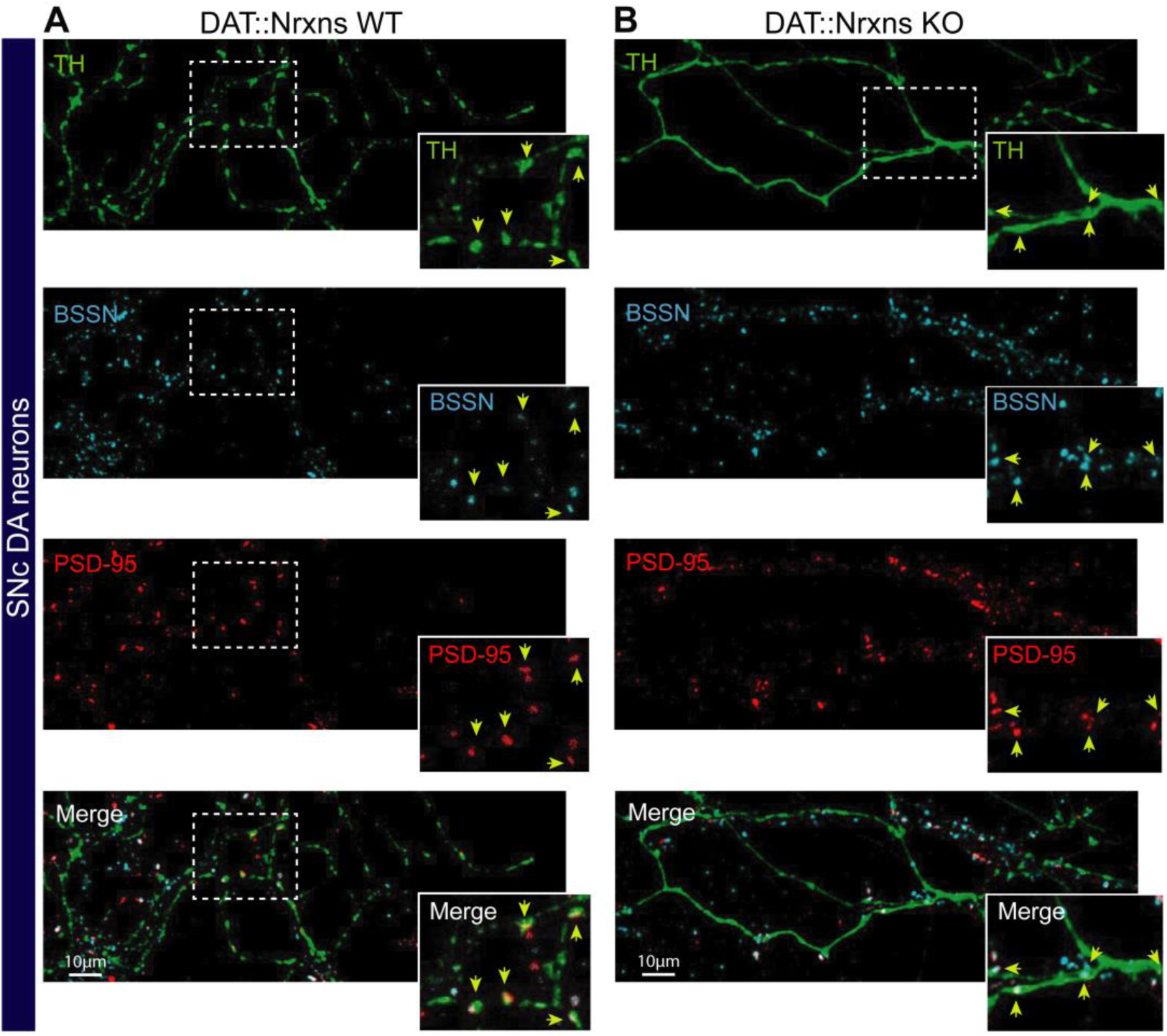
Unchanged proportion of excitatory synapses established by cultured DA neurons after conditional deletion of neurexins. **A** and **B-** Representative images illustrating TH-positive terminals in the axonal arbor of a SNc DA neuron from DAT::NrxnsWT (**A**) and DAT::NrxnsKO (**B**) mice. Bassoon was expressed sparsely and colocalized with the postsynaptic marker PSD95. Yellow arrows indicate regions where TH, Bassoon, and Gephyrin are colocalized.

## Notes

### Competing Interest Statement

The authors have declared no competing interest.

## References

1. Aoto J, Martinelli DC, Malenka RC, Tabuchi K, Südhof TC (2013) Presynaptic neurexin-3 alternative splicing trans-synaptically controls postsynaptic AMPA receptor trafficking. Cell 154:75–88.

2. Banerjee A, Imig C, Balakrishnan K, Kershberg L, Lipstein N, Uronen R-L, Wang J, Cai X, Benseler F, Rhee JS, Cooper BH, Liu C, Wojcik SM, Brose N, Kaeser PS (2022) Molecular and functional architecture of striatal dopamine release sites. Neuron 110:248–265.e9.

3. Banerjee A, Lee J, Nemcova P, Liu C, Kaeser PS (2020) Synaptotagmin-1 is the Ca2+ sensor for fast striatal dopamine release. Elife 9:e58359.

4. Bolte S, Cordelières FP (2006) A guided tour into subcellular colocalization analysis in light microscopy. J Microsc 224:213–232.

5. Boucard AA, Chubykin AA, Comoletti D, Taylor P, Südhof TC (2005) A splice code for trans-synaptic cell adhesion mediated by binding of neuroligin 1 to alpha- and beta-neurexins. Neuron 48:229–236.

6. Boxer EE, Aoto J (2022) Neurexins and their ligands at inhibitory synapses. Front Synaptic Neurosci 14:1087238.

7. Brockhaus J, Kahl I, Ahmad M, Repetto D, Reissner C, Missler M (2024) Conditional Knockout of Neurexins Alters the Contribution of Calcium Channel Subtypes to Presynaptic Ca2+ Influx. Cells 13:981.

8. Bulumulla C, Walpita D, Iyer N, Eddison M, Patel R, Alcor D, Ackerman D, Beyene AG (2024) Synaptic Specializations at Dopamine Release Sites Orchestrate Efficient and Precise Neuromodulatory Signaling. Available at: http://biorxiv.org/lookup/doi/10.1101/2024.09.16.613338 [Accessed October 29, 2025].

9. Burke S, Trudeau L-E (2022) Axonal Domain Structure as a Putative Identifier of Neuron-Specific Vulnerability to Oxidative Stress in Cultured Neurons. eNeuro 9:ENEURO.0139-22.2022.

10. Chen F, Venugopal V, Murray B, Rudenko G (2011) The structure of neurexin 1α reveals features promoting a role as synaptic organizer. Structure 19:779–789.

11. Cheung A, Konno K, Imamura Y, Matsui A, Abe M, Sakimura K, Sasaoka T, Uemura T, Watanabe M, Futai K (2023) Neurexins in serotonergic neurons regulate neuronal survival, serotonin transmission, and complex mouse behaviors. Elife 12:e85058.

12. Chih B, Gollan L, Scheiffele P (2006) Alternative Splicing Controls Selective Trans-Synaptic Interactions of the Neuroligin-Neurexin Complex. Neuron 51:171–178.

13. Craig AM, Banker G, Chang W, McGrath ME, Serpinskaya AS (1996) Clustering of gephyrin at GABAergic but not glutamatergic synapses in cultured rat hippocampal neurons. J Neurosci 16:3166–3177.

14. Dal Bo G, St-Gelais F, Danik M, Williams S, Cotton M, Trudeau L-E (2004) Dopamine neurons in culture express VGLUT2 explaining their capacity to release glutamate at synapses in addition to dopamine. J Neurochem 88:1398–1405.

15. Dean C, Scholl FG, Choih J, DeMaria S, Berger J, Isacoff E, Scheiffele P (2003) Neurexin mediates the assembly of presynaptic terminals. Nat Neurosci 6:708–716.

16. Delignat-Lavaud B, Kano J, Ducrot C, Massé I, Mukherjee S, Giguère N, Moquin L, Lévesque C, Burke S, Denis R, Bourque M-J, Tchung A, Rosa-Neto P, Lévesque D, De Beaumont L, Trudeau L-É (2023) Synaptotagmin-1-dependent phasic axonal dopamine release is dispensable for basic motor behaviors in mice. Nat Commun 14:4120.

17. Descarries L, Bosler O, Berthelet F, Des Rosiers MH (1980) Dopaminergic nerve endings visualised by high-resolution autoradiography in adult rat neostriatum. Nature 284:620–622.

18. Ducrot C, Bourque M-J, Delmas CVL, Racine A-S, Guadarrama Bello D, Delignat-Lavaud B, Domenic Lycas M, Fallon A, Michaud-Tardif C, Burke Nanni S, Herborg F, Gether U, Nanci A, Takahashi H, Parent M, Trudeau L-E (2021) Dopaminergic neurons establish a distinctive axonal arbor with a majority of non-synaptic terminals. FASEB J 35:e21791.

19. Ducrot C, de Carvalho G, Delignat-Lavaud B, Delmas CVL, Halder P, Giguère N, Pacelli C, Mukherjee S, Bourque M-J, Parent M, Chen LY, Trudeau L-E (2023) Conditional deletion of neurexins dysregulates neurotransmission from dopamine neurons. Elife 12:e87902.

20. Fasano C, Thibault D, Trudeau L-E (2008) Culture of postnatal mesencephalic dopamine neurons on an astrocyte monolayer. Curr Protoc Neurosci Chapter 3:Unit 3.21.

21. Fulton S, Thibault D, Mendez JA, Lahaie N, Tirotta E, Borrelli E, Bouvier M, Tempel BL, Trudeau L-E (2011) Contribution of Kv1.2 voltage-gated potassium channel to D2 autoreceptor regulation of axonal dopamine overflow. J Biol Chem 286:9360–9372.

22. Giguère N, Delignat-Lavaud B, Herborg F, Voisin A, Li Y, Jacquemet V, Anand-Srivastava M, Gether U, Giros B, Trudeau L-É (2019) Increased vulnerability of nigral dopamine neurons after expansion of their axonal arborization size through D2 dopamine receptor conditional knockout. PLoS Genet 15:e1008352.

23. Graf ER, Zhang X, Jin S-X, Linhoff MW, Craig AM (2004) Neurexins induce differentiation of GABA and glutamate postsynaptic specializations via neuroligins. Cell 119:1013–1026.

24. Ichtchenko K, Hata Y, Nguyen T, Ullrich B, Missler M, Moomaw C, Südhof TC (1995) Neuroligin 1: a splice site-specific ligand for beta-neurexins. Cell 81:435–443.

25. Kornau HC, Seeburg PH, Kennedy MB (1997) Interaction of ion channels and receptors with PDZ domain proteins. Curr Opin Neurobiol 7:368–373.

26. Liu C, Kershberg L, Wang J, Schneeberger S, Kaeser PS (2018) Dopamine Secretion Is Mediated by Sparse Active Zone-like Release Sites. Cell 172:706–718.e15.

27. Love MI, Huber W, Anders S (2014) Moderated estimation of fold change and dispersion for RNA-seq data with DESeq2. Genome Biol 15:550.

28. Luo F, Sclip A, Jiang M, Südhof TC (2020) Neurexins cluster Ca2+ channels within the presynaptic active zone. EMBO J 39:e103208.

29. Matsuda W, Furuta T, Nakamura KC, Hioki H, Fujiyama F, Arai R, Kaneko T (2009) Single nigrostriatal dopaminergic neurons form widely spread and highly dense axonal arborizations in the neostriatum. J Neurosci 29:444–453.

30. Mendez JA, Bourque M-J, Dal Bo G, Bourdeau ML, Danik M, Williams S, Lacaille J-C, Trudeau L-E (2008) Developmental and target-dependent regulation of vesicular glutamate transporter expression by dopamine neurons. J Neurosci 28:6309–6318.

31. Mendez JA, Bourque M-J, Fasano C, Kortleven C, Trudeau L-E (2011) Somatodendritic dopamine release requires synaptotagmin 4 and 7 and the participation of voltage-gated calcium channels. J Biol Chem 286:23928–23937.

32. Missler M, Zhang W, Rohlmann A, Kattenstroth G, Hammer RE, Gottmann K, Südhof TC (2003) Alpha-neurexins couple Ca2+ channels to synaptic vesicle exocytosis. Nature 423:939–948.

33. Pacelli C, Giguère N, Bourque M-J, Lévesque M, Slack RS, Trudeau L-É (2015) Elevated Mitochondrial Bioenergetics and Axonal Arborization Size Are Key Contributors to the Vulnerability of Dopamine Neurons. Current Biology 25:2349–2360.

34. Parent M, Parent A (2006) Relationship between axonal collateralization and neuronal degeneration in basal ganglia. J Neural Transm Suppl:85–88.

35. Schultz W (2007) Multiple dopamine functions at different time courses. Annu Rev Neurosci 30:259–288.

36. Stuber GD, Hnasko TS, Britt JP, Edwards RH, Bonci A (2010) Dopaminergic terminals in the nucleus accumbens but not the dorsal striatum corelease glutamate. J Neurosci 30:8229–8233.

37. Südhof TC (2017) Synaptic Neurexin Complexes: A Molecular Code for the Logic of Neural Circuits. Cell 171:745–769.

38. Sulzer D, Joyce MP, Lin L, Geldwert D, Haber SN, Hattori T, Rayport S (1998) Dopamine neurons make glutamatergic synapses in vitro. J Neurosci 18:4588–4602.

39. Tritsch NX, Ding JB, Sabatini BL (2012) Dopaminergic neurons inhibit striatal output through non-canonical release of GABA. Nature 490:262–266.

40. Tritsch NX, Granger AJ, Sabatini BL (2016) Mechanisms and functions of GABA co-release. Nat Rev Neurosci 17:139–145.

41. Uchigashima M, Cheung A, Suh J, Watanabe M, Futai K (2019) Differential expression of neurexin genes in the mouse brain. J Comp Neurol 527:1940–1965.

42. Uchigashima M, Ohtsuka T, Kobayashi K, Watanabe M (2016) Dopamine synapse is a neuroligin-2-mediated contact between dopaminergic presynaptic and GABAergic postsynaptic structures. Proc Natl Acad Sci U S A 113:4206–4211.

43. Ullrich B, Ushkaryov YA, Südhof TC (1995) Cartography of neurexins: more than 1000 isoforms generated by alternative splicing and expressed in distinct subsets of neurons. Neuron 14:497–507.

44. Ushkaryov YA, Petrenko AG, Geppert M, Südhof TC (1992) Neurexins: Synaptic Cell Surface Proteins Related to the α-Latrotoxin Receptor and Laminin. Science 257:50–56.

45. Wang A, Xiang Y-Y, Yang BB, Lu W-Y (2019) Neurexin-1α regulates neurite growth of rat hippocampal neurons. Int J Physiol Pathophysiol Pharmacol 11:115–125.

46. Zhang C, Atasoy D, Araç D, Yang X, Fucillo MV, Robison AJ, Ko J, Brunger AT, Südhof TC (2010) Neurexins physically and functionally interact with GABA(A) receptors. Neuron 66:403–416.

